# Peripatric speciation associated with genome expansion and female-biased sex ratios in the moss genus *Ceratodon*

**DOI:** 10.1101/227280

**Authors:** Marta Nieto-Lugilde, Olaf Werner, Stuart F. McDaniel, Petr Koutecký, Jan Kučera, Samah Mohamed Rizk, Rosa M. Ros

**Affiliations:** Departamento de Biología Vegetal, Facultad de Biología, Universidad de Murcia, Campus de Espinardo, 30100 Murcia, Spain.; Biology Department, University of Florida, Gainesville, Florida 32611, USA.; Faculty of Science, University of South Bohemia, Branišovská 31, CZ-370 05 České Budějovice, Czech Republic.; Genetics Department, Faculty of Agriculture, Ain Shams University, 68 Hadayek Shubra, 11241 Cairo, Egypt.

**Keywords:** Bryophyta, cosmopolitan species, DNA sequencing, flow cytometry, hybridization, Mediterranean mountains, phylogenetic data, polyploidy.

## Abstract

**PREMISE OF THE STUDY:** A period of allopatry is widely believed to be essential for the evolution of reproductive isolation. However, strict allopatry may be difficult to achieve in some cosmopolitan, spore-dispersed groups, like mosses. Here we examine the genetic and genome size diversity in Mediterranean populations of the moss *Ceratodon purpureus* s.l. to evaluate the role of allopatry and ploidy change in population divergence.

**METHODS:** We sampled populations of the genus *Ceratodon* from mountainous areas and lowlands of the Mediterranean region, and from western and central Europe. We performed phylogenetic and coalescent analyses on sequences from five nuclear introns and a chloroplast locus to reconstruct their evolutionary history. We also estimated the genome size using flow cytometry, employing propidium iodide, and determined their sex using a sex-linked PCR marker.

**KEY RESULTS:** Two well differentiated clades were resolved, discriminating two homogeneous groups: the widespread *C. purpureus* and a local group mostly restricted to the mountains in southern Spain. The latter also possessed a genome size 25% larger than the widespread *C. purpureus*, and the samples of this group consist entirely of females. We also found hybrids, and some of them had a genome size equivalent to the sum of the *C. purpureus* and Spanish genome, suggesting that they arose by allopolyploidy.

**CONCLUSIONS:** These data suggest that a new species of *Ceratodon* arose via peripatric speciation, potentially involving a genome size change and a strong female-biased sex ratio. The new species has hybridized in the past with *C. purpureus*.

## INTRODUCTION

The origin of new species represents a major unsolved problem in evolutionary biology (Rieseberg and Willis, 2007; Seehausen et al., 2014; Dev, 2015). Theory shows that the simplest mechanism for generating new species is through allopatric speciation, in which some portion of a species’ range becomes geographically isolated, allowing natural selection or genetic drift to drive allele frequency changes that ultimately generate additional reproductive barriers (Mayr, 1963; Barraclough and Vogler, 2000; Coyne and Orr, 2004). This is because even modest levels of gene flow can homogenize allele frequencies between populations, retarding divergence (Wright, 1931). While local adaptation can drive peripatric or sympatric divergence in cases where the immigrant rate is less than the intensity of selection (Lenormand, 2002), most empirical studies cannot exclude the possibility that speciation was preceded by a period of allopatry (Nadachowska-Brzyska et al., 2013; Shaner et al., 2015). This presents a paradox in species-rich groups like mosses, where long-distance migration appears to be common, but speciation and diversification have occurred in spite of the fact that geographic barriers may not cause a long-term impediment to gene flow (Shaw et al., 2003; Piñeiro et al., 2012; Lewis et al., 2014a; Szövényi et al., 2014; Barbé et al., 2016).

One potential resolution to this paradox is sympatric speciation through polyploidy, which is frequent in flowering plants (Ramsey and Schemske, 1998; Mallet, 2005), and potentially in mosses (Crosby, 1980; Kuta and Przywara, 1997; Såstad, 2005; McDaniel et al., 2010; Rensing et al., 2013). Polyploidy generates a strong reproductive barrier in a single mutational event (Ramsey and Schemske, 1998; Madlung, 2013). However, the homogeneity in bryophyte genome sizes (Voglmayr, 2000) raises the possibility that the role played by polyploidy in moss speciation may be small relative to other speciation mechanisms. The nature of the genomic, demographic, or ecological factors beyond geographic isolation and polyploidy that generate reproductive barriers between nascent species of mosses remain poorly characterized (McDaniel et al., 2010; Yousefi et al., 2017).

Within mosses, the genetic basis of reproductive barriers is best characterized among populations of *Ceratodon purpureus* (Hedw.) Brid. (Ditrichaceae) (McDaniel et al., 2007, 2008). Moreover, the developing genomic and laboratory tools make this species a promising model for further ecological genomic study (McDaniel et al., 2016). *Ceratodon purpureus* is abundant on every continent, and grows on wide variety of substrates (Crum, 1973). Molecular population genetic analyses indicated that gene flow among northern and even southern hemisphere populations was frequent but tropical populations were more genetically isolated (McDaniel and Shaw, 2005). These observations suggest that the current level of sampling may be insufficient to detect the full scope of population structure among populations in this taxon. Indeed, partial hybrid breakdown was clearly evident in crosses between a temperate and a tropical population, suggesting that reproductive barriers may be in the process of evolving between ecologically distinct regions of the distribution of *C. purpureus* (McDaniel et al., 2007, 2008). These barriers did not involve ploidy differences. However, the genome size of *C. purpureus* is well-characterized in only a modest number of European samples (0.39 pg s.d. 0.0046, n=10, Voglmayr, 2000), leaving open the possibility that polyploidy contributes to reproductive isolation among isolates from other parts of its broad cosmopolitan distribution.

In a previous phylogeographic analysis (McDaniel and Shaw, 2005), the Mediterranean region contained several rare haplotypes that were distantly related to the common haplotypes found throughout the range of *C. purpureus*. Here we sought to test for the existence of any relationship between the genetic diversity and DNA content found in the Mediterranean area in the moss genus *Ceratodon*. McDaniel and Shaw (2005) argued that frequent gene flow maintained the genetic homogeneity of the species, at least among the temperate Northern Hemisphere populations, but that the divergent populations were simply outside the main area of spore rain, and therefore had not yet been homogenized. Alternatively, these isolated populations could represent cryptic species, and reproductive isolation evolved in spite of this gene flow (McDaniel et al., 2007, 2008). To distinguish between these alternatives, we evaluated the patterns of polymorphism in five nuclear introns and a single chloroplast locus in plants sampled from mountainous areas of the Mediterranean region and other mountain regions and lowlands mostly from southern Europe. We also estimated the genome size of these isolates using flow cytometry. These data clearly show that a new species have evolved within the genus *Ceratodon*, accompanied by both large non-polyploid and allopolyploid changes in genome size, and potentially major changes in sexual system. These insights also highlight the complexity of peripatric speciation mechanisms in bryophytes.

## MATERIALS AND METHODS

### Plant material

For this study we generated genetic data for a total of 93 samples, 71 (76.4%) from Mediterranean mountain areas (47 from Spanish Sierra Nevada Mountains, 19 from Spanish central mountain ranges, three from Spanish south-eastern mountains, and two from Sicilian Mount Etna). Of the remaining 22 samples, 11 (11.8%) were from other European mountainous systems (eight from the Alps and three from the Pyrenees) and 11 specimens (11.8%) were from lowlands (three from Czech Republic, two from Germany, two from Sweden, two from United Kingdom, and two from South Africa). Mainly between April and November 2011-2014 (more detailed information in Appendix 1) we collected 84 new samples for this study, all of which are deposited at MUB (Herbarium of the University of Murcia, Spain), nine samples were loaned from herbaria, including BOL (Bolus Herbarium, University of Cape Town, South Africa), CBFS (University of South Bohemia, Czech Republic), S (Herbarium of the Swedish Museum of Natural History, Sweden), and two samples were donated from Laura Forrest (at Royal Botanic Garden Edinburgh, United Kingdom). We sequenced four specimens of *Cheilothela chloropus* (Brid.) Lindb. to use as an outgroup (Voucher information and Genbank accession numbers are listed in Appendix 1).

### DNA sequencing

To examine the genealogical relationships among the 93 isolates, we sequenced five nuclear exon-primed intron-spanning loci, including *rpL23A* and *TRc1b3.05* (McDaniel et al., 2013a; referenced by EST accessions AW086590 and AW098560), *hp23.9*, *PPR* and *TBP* (McDaniel et al., 2013a, b), and a single chloroplast locus (*trn*L). We amplified all loci from all individuals in 20 μL polymerase chain reaction using Thermo Scientific DreamTaq DNA Polymerase (Thermo Fisher Scientific Inc.). The cycling conditions were 94°C for 2 min, then 10 cycles of 94°C for 15 s, an annealing temperature of 65°C that dropped one degree each cycle, and 72°C for 1 min, followed by 20 cycles of 94°C for 15 s, 56°C for 30 s, and 72°C for 1 min, and terminating with 72°C for 7 min (McDaniel et al., 2013b). The resulting PCR products were ready to use for sequencing removing unincorporated primers and inactivates unincorporated nucleotides using PCR clean-up reaction with Exo I (Thermo Fisher Scientific Inc.) and FastAP Alkaline Phosphatase enzymes (Thermo Fisher Scientific Inc.). Both enzymes were heat-inactivated by maintaining the mixture at 85°C for 15 min. Sequencing was accomplished on an ABI3730XL DNA Analyzer, Applied Biosystems (Macrogen Europe, The Netherlands, Amsterdam).

### Cloning of DNA sequences

In samples where we observed double peaks in the chromatograms, we cloned all loci. PCR products were isolated from agarose gels, and cloned using the CloneJet PCR Cloning Kit (ThermoFisher Scientific, Spain). Cloning efficiency and accuracy were checked using PCR reactions. Successful clones then were sequenced using an ABI3730XL DNA Analyzer (Macrogen).

### Phylogenetic analyses

We aligned the DNA sequences using CLUSTALW (Larkin et al., 2007) as implemented in Bioedit (Hall, 1999) and manually resolved inconsistencies in the resulting alignment. DnaSP v5 (Librado and Rozas, 2009) was used to observe characteristics such as total length with and without gaps, number of constant positions and number of parsimony-informative variable positions about all loci. We coded gaps as informative with a simple indel coding strategy (Simmons and Ochoterena, 2000) implemented in SeqState (Müller, 2005). We performed phylogenetic analyses using MrBayes v.3.2 (Ronquist et al., 2012). The need for a priori model testing was removed using the substitution model space in the Bayesian MCMC analysis itself (Huelsenbeck et al., 2004) with the option nst=mixed. The sequence and indel data were treated as separate and unlinked partitions. The a priori probabilities supplied were those specified in the default settings of the program. Posterior probability distributions of trees were generated using the Metropolis-coupled Markov chain Monte Carlo (MCMCMC) method. To search for convergence in the phylogenetic analyses we used two runs with different setting for some of the loci. For *hp23.9*, *TBP* and *trn*L, four chains with 1 x 10^7^ generations were run simultaneously, with the temperature of the single heated chain set to the default in MrBayes. Nevertheless eight chains with 1 x 10^6^ generations each one were run, changing the temperature of the single heated chain set to 2 (*PPR*), 3 (*TRc1b3.05*) and 6 (*rpL23A*), because with the default temperature setting convergence was not reached in initial runs. Chains were sampled every 1000 generations and the respective trees were written into a tree file. The first 25% of the total sampled trees of each run were discarded as burnin. Consensus trees and posterior probabilities of clades were calculated by combining the two runs and using the trees sampled after the chains converged and had become stationary. The sump command of MrBayes was used to check whether an appropriate sample from the posterior was obtained. To do so, we first inspected visually the log likelihood plot, which should not show tendencies to decrease or increase over time and the different runs should show similar values. Then we checked that the effective sampling size (ESS) values for all parameters reached at least 500 and that the Potential Scale Reduction Factor (PSRF) for each parameter was close to 1.00. The genealogies were rooted with sequences from *Cheilothela chloropus.* The final trees were edited with TreeGraph2 (Stöver and Müller, 2010). We performed phylogenetic analyses using the same setting as before combining the new sequences generated here with other sequences for the *TBP* locus available on GenBank from Antarctica (1), Australia (1), and Eastern North America (54), which were previously reported by McDaniel et al. (2013b).

Low resolution in phylogenetic reconstructions can sometimes be caused by incongruence or conflicts in the molecular datasets that lead to different equally possible solutions (Huson and Bryant, 2006; Draper et al., 2015). To evaluate this possibility, we reconstructed a phylogenetic network based on the neighbor-net method (Bryant and Moulton, 2004) using the program SplitsTree4, version 4.13.1 (Huson and Bryant, 2006) for the six concatenated loci. The calculations were based on uncorrected p-distances. To test the hypothesis of recombination in each graph, a pairwise homoplasy index (Phi-test) was calculated, which is a robust and reliable statistic to detect recombination. This estimates the mean refined incompatibility score from nearby sites. Under the null hypothesis of no recombination, the genealogical correlation of adjacent sites is invariant to permutations of the sites as all sites have the same history. In the case of finite levels of recombination, the order of the sites is important, as distant sites will tend to have less genealogical correlation than adjacent sites (Bruen et al., 2006). The significance is then tested using a permutation test by default. In accordance with Bruen et al. (2006) for the Phi test of recombination, p-value < 0.05 indicates the presence of recombination signal.

### Coalescent stochasticity analyses

Individual gene trees often differ from each other and from the species tree (Rosenberg, 2002; Mao et al., 2014). To check whether the differentiation we found between the Sierra Nevada (SN) and Worldwide (Ww) clades was a good fit to the multispecies coalescent model (MSCM), we employed an approach based on posterior predictive simulation implemented in the R statistical language package, “P2C2M” (Gruenstaeudl et al., 2016). In this approach, a posterior distribution of gene genealogies estimated from empirical data is compared to a posterior predictive distribution of genealogies simulated under a model of interest. We first used *BEAST (Heled and Drummond, 2010) to infer genealogies and species trees, and the simulation of genealogies under the MSCM with ms program (Hudson, 2002) under a JC +I +G nucleotide substitution model selected as the most probable in all loci using jModelTest (Posada, 2008). Gaps were included as a character state in these analyses. The run was conducted assuming a strict clock for each locus. We selected “Yule Model” as the species tree prior, and employed “Piecewise linear and constant root” as the population size model. Finally, the default values for MCMC analysis were used. We compared the genealogies from the posterior distribution to the species trees and from the posterior predictive distribution to the species tree using two descriptive summary statistics: *lcwt* (likelihood of the coalescent waiting times) and the number of deep coalescences (*ndc*). When samples are drawn from data with a good fit to the MSCM, the summary statistics from each distribution should be approximately equal and the expected difference between the two is zero (Reid et al., 2014). Data that are a poor fit to the MSCM are indicted by a deviation from the expectation of a difference distribution that is centered on zero is encountered above a specified quantile level (Gruenstaeudl et al., 2016).

### Genome size determination

We used flow cytometry (FCM) technology for 75 specimens to estimate nuclear DNA content. One shoot of each sample was chopped with a razor blade together with the internal standard *Carex acutiformis* Ehrh. 1C = 0.41 pg, Lipnerová et al., 2012) or *Bellis perennis* L. (1C = 1.56 pg; our own calibration against *Carex acutiformis*) in 1 ml of LB01 buffer (Doležel et al., 1989). The fluorochrome propidium iodide and RNase IIa (both at final concentration 50 μg/ml) were added immediately; the samples were stained for at least 10 minutes. The samples were analyzed using a Partec CyFlow SL flow cytometer equipped with a 532 nm (green) diode-pumped solid-state laser (100 mW output); the fluorescence intensity of 12000 particles was recorded. When possible, we used in vitro cultivated fresh material, but for 47 samples that did not grow satisfactorily in vitro, we used dry material collected in the years 2009-2014. The fluorescence histograms were processed using FlowJo v 10.2 software (TreeStar Inc.).

### Sex determination

To determinate sex, one plant per sample was employed. We amplified the *rpS15A* sex-linked locus by PCR and digested the product with HindIII. An intron in the *rpS15A* amplicon contains a cut-site difference between the male and female products (Norrell et al., 2014) which is clearly observable in the banding patterns which were visualized after electrophoresis in an agarose gel and scored by hand. We identified the sex of 82 samples, 88.17 % of the total, which were from Sierra Nevada Mountains (42), Spanish central mountain ranges (16), Spanish south-eastern mountains (3), Sicilian Mount Etna (2), Alps (7), Pyrenees (3), South Africa (2), Germany (2), Czech Republic (3), and Sweden (2). For the remaining samples we could not unambiguously interpret the pattern in the restriction-site fragment length polymorphism in the *rpS15A* amplicon. We express the results as a proportion of males and computed the 95% confidence interval for this estimate with the *dbinom* function in R (R Development Core Team, 2017).

### Calculation of the binomial proportion confidence interval

If the total number of experiments and the number of positive outcomes of a success-failure experiment are known, it is possible to calculate the confidence intervals (CI) for the probability of success. As a consequence, the CI for the proportion of males (or females) in a population (success) based on the results of sex-determination of a given number of individuals can be calculated. We used the Hmisc package (Harrell, 2018) in R3.4.3 (R Development Core Team, 2017), with the options “Wilson” and “Exact” (=Clopper-Pearson) to calculate the CI.

## RESULTS

### Phylogenetic analyses

The sequence alignments varied in total length between 207 (215 with coded gaps) to 848 (891) positions, for *hp23.9* and *rpL23A*, respectively. The number of constant positions was between 186 and 715 for the above mentioned loci and the parsimony-informative variable positions differed between 5 and 95 for *trn*L and *rpL23A*, respectively (Table 1). The loci *PPR, TBP, rpL23A*, and *TRc1b3.05* showed two well differentiated clades with support of 0.87 (0.85-0.88)-0.77 (0.75-0.79) posterior probability (pp), 0.96 (0.95-0.96)-1.00 (1.00-1.00) pp, 1.00 (1.00-1.00)-1.00 (1.00-1.00) pp, and 1.00 (1.00-1.00)-1.00 (1.00-1.00) pp, respectively (Fig. 1, see Supplemental Data with this article, Appendices S1, S2, S3). In the case of *rpL23A*, sequences of *Cheilothela chloropus* were not obtained for use as outgroup, but again two clades were resolved. The *hp23.9* locus had a support for one clade of 1.00 (1.00-1.00) pp but the other clade had a value of 0.55 (0.45-0.61) pp (Appendix S4). In all the five nuclear loci studied, one of the clades was formed always by 34 Sierra Nevada Mountains samples and one of the Spanish south-eastern mountains; we refer to this as the SN group. The second clade consistently included 42 specimens coming from the rest of the sampled areas, including one from Sierra Nevada and two from Spanish south-eastern mountains; we refer to this as the Ww group. For one marker (*TBP*) we added sequences available at GenBank, including samples from Antarctica, Australia, and North America. The resulting tree topology shows that our samples give a reasonable good representation of the Ww group and that none of these additional sequences is closely related to the SN samples (Appendix S5). The remaining 17 sequenced samples were strongly resolved in either the SN clade or the Ww clade, depending on the studied locus (they did not present intermediate sequences between both clades, Appendix S6); we considered these samples recombinants. The term “hybrid” applied to bryophytes should strictly be used only for the sporophytic hybrids (2n) (Anderson, 1980); for their gametophytic progeny (n) showing combination of parental alleles after meiosis “recombinants” should be used (Shaw, 1994, 1998) in order not to confuse with hybrids observed among vascular plants. The recombinants derived mainly from SN Mountains, but also from Spanish central mountain ranges, the Alps and the lowlands of the United Kingdom (Fig. 2). The chloroplast locus showed one well supported clade 0.96 (0.95-0.98) pp, and all remaining samples with deeper coalescent events (Fig. 1). All the samples considered as recombinants based on the nuclear markers were closely related and sister to the rest of the SN samples, with the only exception of one specimen from Sierra Nevada Mountains (MUB 49528), which is a recombinant and belongs to the Ww chloroplast clade.

**Table 1.**
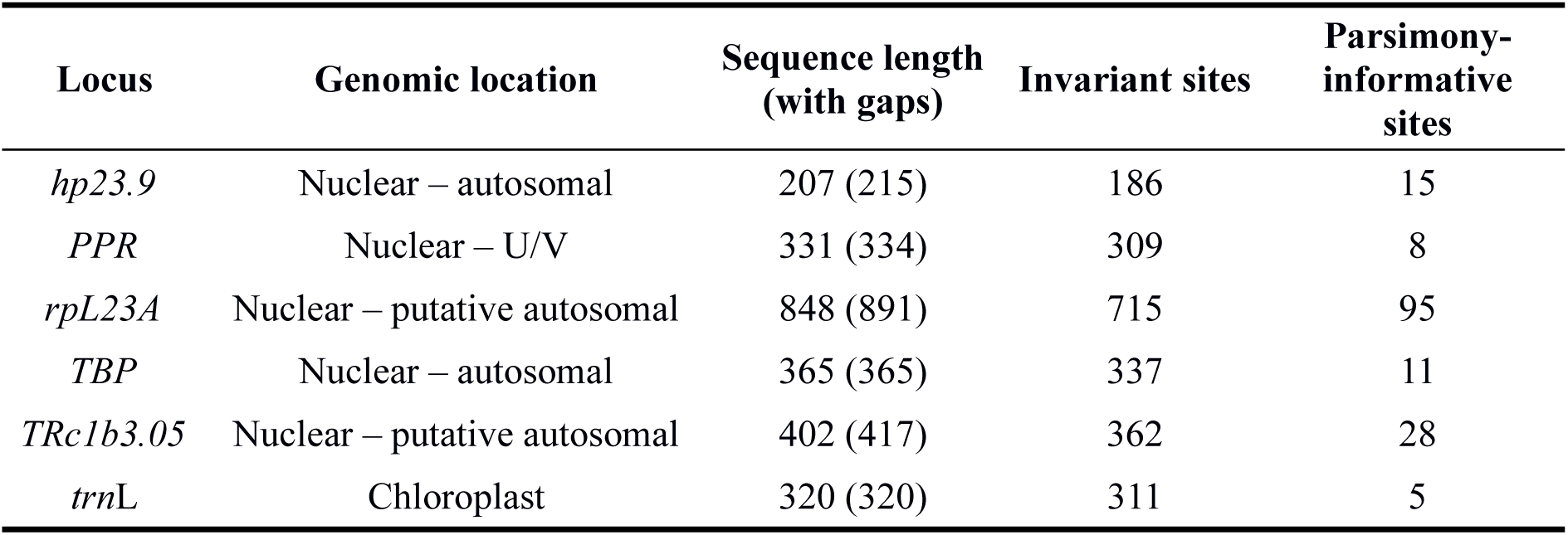
Characteristics of the loci used for molecular evolutionary analyses. The genomic location “nuclear - putative autosomal” is based on unpublished data.

| Locus | Genomic location | Sequence length<br>(with gaps) | Invariant sites | Parsimony-informative<br>sites |
| --- | --- | --- | --- | --- |
| *hp23.9* | Nuclear – autosomal | 207 (215) | 186 | 15 |
| *PPR* | Nuclear – U/V | 331 (334) | 309 | 8 |
| *rpL23A* | Nuclear – putative autosomal | 848 (891) | 715 | 95 |
| *TBP* | Nuclear – autosomal | 365 (365) | 337 | 11 |
| *TRc1b3.05* | Nuclear – putative autosomal | 402 (417) | 362 | 28 |
| *trnL* | Chloroplast | 320 (320) | 311 | 5 |

**Fig. 1.**
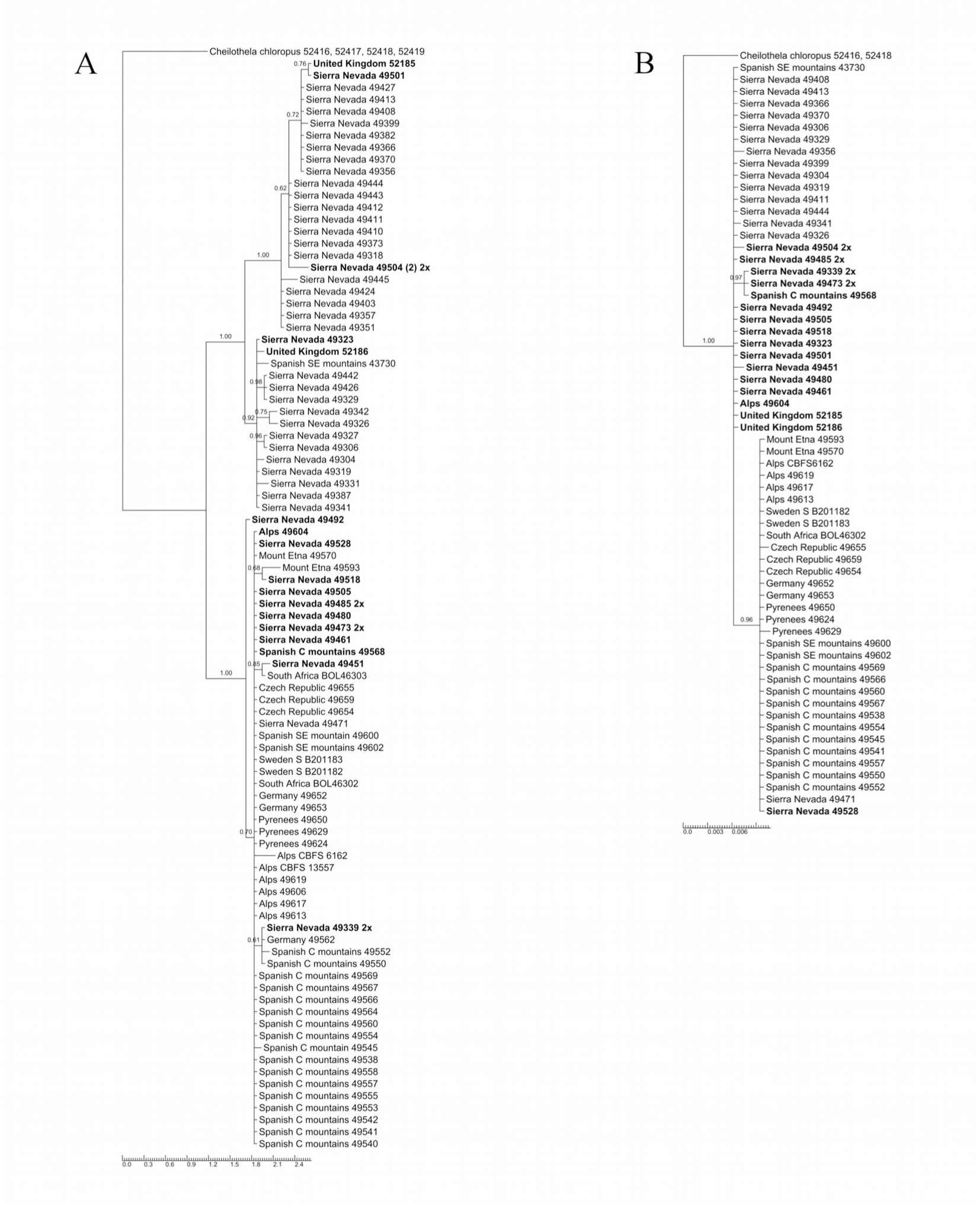
Phylogenetic trees inferred from two of the studied loci. For each tip in the trees geographical origin and number of herbarium are given (numbers without letters are from MUB); 2x is used to highlight diploid samples; number of equal sequences obtained by cloning is indicated between parentheses if there was more than one; bold letters indicate recombinant samples. A) From nuclear *TRc1b3.05* locus and B) From chloroplast *trn*L locus.

**Fig. 2.**
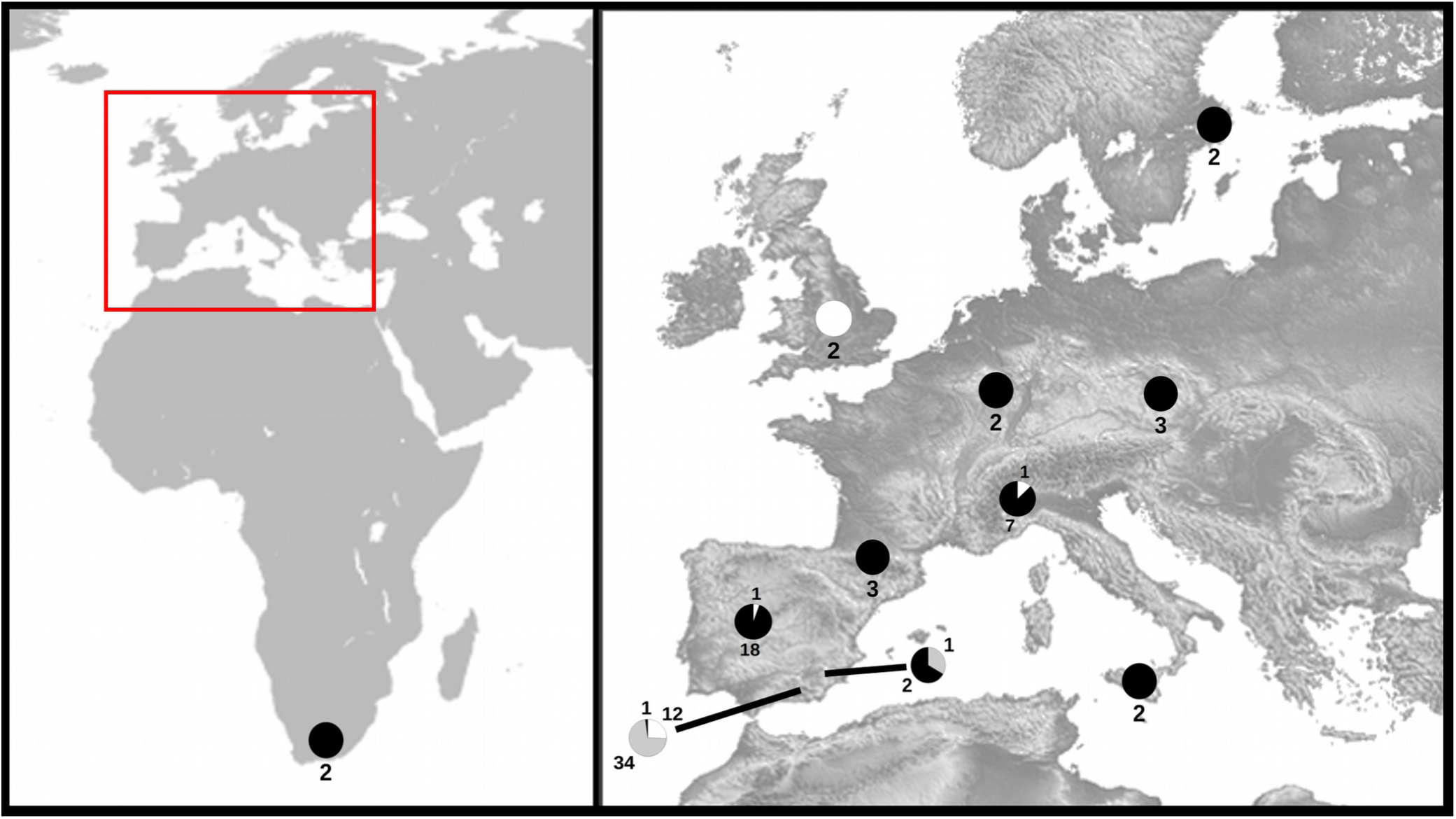
Geographic location of *Ceratodon* samples included in this study. Pie charts indicate proportion of samples of each genomic group by areas (black: Ww genome group; grey: SN genome group; white: recombinant samples). The number of samples by groups in each area is given.

**Fig. 3.**
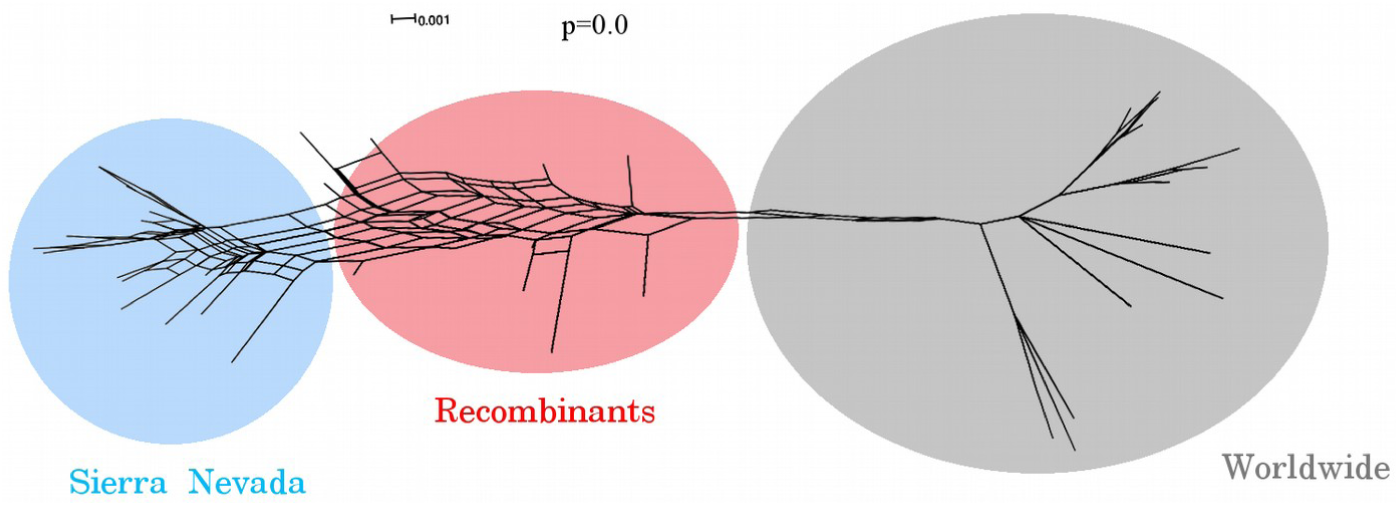
Neighbor-Net network to test signals of reticulate evolution between the samples. The main groups are highlight by color circles with its names. The p value from the Phi test of recombination is indicated.

**Fig. 4.**
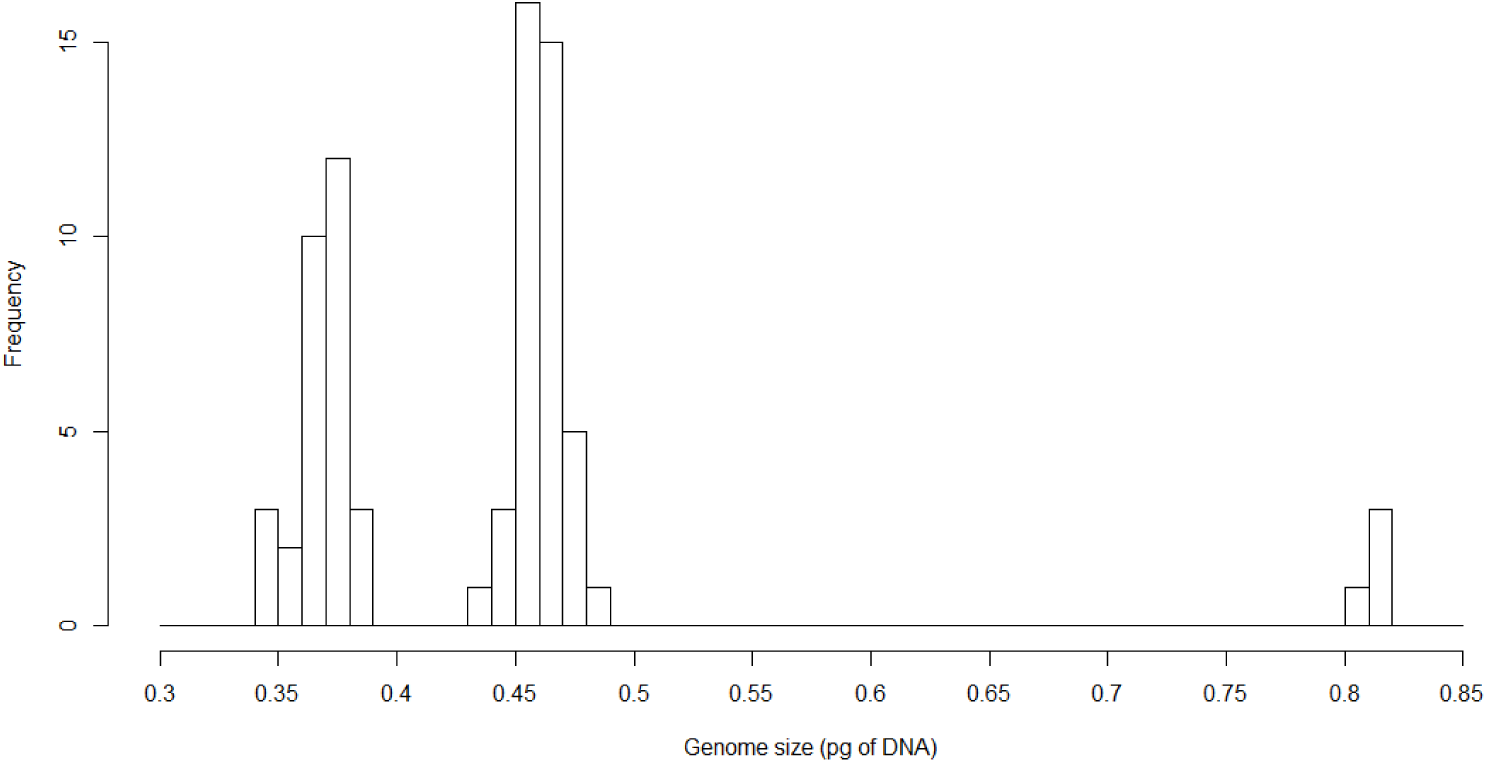
Histogram of genome sizes of representative samples of *Ceratodon* generated by flow cytometry. A conversion factor of 0.85 was applied to the data obtained from dry material.

The apparent uncertain position of some individuals is clarified by the result of the Neighbor-Net network (Fig. 3). Moreover for the phi-test when the six loci were studied together, a highly significant p-value (0.0) was obtained, confirming the presence of recombination signal. Graphically, two extreme groups can be observed, the SN group and the Ww group, with some individuals in intermediate positions.

### Cloning DNA sequences

Loci cloning confirmed that diploid specimens (see Flow cytometry analyses results) present two different copies of the same loci in most cases. The loci *TRc1b3.05*, *PPR* and *rpL23A* presented predominantly a single copy, although some individuals presented the two copies in other loci (Appendix S6). Some haploid individuals presented two different copies of a locus. This may be due to the possibility of gene redundancy, which can result from unequal crossing over, retroposition or chromosomal (or genome) duplication (Magadum et al., 2013).

### Coalescent stochasticity analyses

Although our data suggested the existence of recombinants between the two groups, incomplete lineage sorting and hybridization may result in similar molecular signals. Nevertheless, the two summary statistics *ndc* and *lcwt* employed for the comparison of the genealogies from the posterior distribution to the species trees and from the posterior predictive distribution to the species trees, show significant (p < 0.05) differences between our data with respect to MSCM (Table 2), indicating that incomplete lineage sorting (coalescent model) alone cannot explain the different tree topologies.

**Table 2.** Results of P2C2M analysis in which *lcwt* and *ndc* descriptive summary statistics are shown for each DNA locus analyzed. All loci under study are of nuclear origin, except *trn*L. Asterisks indicate cases of poor model fit at a probability level of 0.05, and. n.a., not applicable.

| Locus | *lcwt* | *ndc* |
| --- | --- | --- |
| *hp23.9* | 510.16 ($\pm 117.86$)* | -57.46 ($\pm 29.93$)* |
| *PPR* | 530.67 ($\pm 121.31$)* | -59.88 ($\pm 29.87$)* |
| *rpL23A* | 462.75 ($\pm 110.74$)* | -53.04 ($\pm 29.16$)* |
| *TBP* | 525.94 ($\pm 120.55$)* | -59.56 ($\pm 29.77$)* |
| *TRc1b3.05* | 516.43 ($\pm 121.96$)* | -58.31 ($\pm 29.96$)* |
| *trnL* | n.a. | -65.33 ($\pm 29.26$)* |
| Sum of all genes | n.a. | -353.59 ($\pm 105.62$)* |

### Flow cytometry analyses

We obtained three clearly differentiated groups of cytotypes for both fresh and dry material (Table 3, Fig. 4). Measurements from dry material gave higher values (by 18% on average) than those from fresh material, for this reason a conversion factor (1/1.18 = 0.85) was employed to the former. When fresh and dry materials are considered together, the first cytotype had a mean value of 1C = 0.37 pg, and the second one showed 25.4% more of DNA content (1C = 0.46 pg). The third cytotype had 1C = 0.82 pg mean value of DNA content. All of specimens of Ww group belonged to the smallest cytotype while those of the SN group were categorized in the second cytotype, and the recombinant specimens were found in both the second and the third cytotype (Appendix S6).

**Table 3.** Nuclear DNA content expressed in pg as measured by flow cytometry. Cytotypes considered, number of samples used in the analyses (N), mean value of DNA, standard deviation and range of values obtained for each cytotype are given (* conversion factor of 0.85 is applied to dry material when fresh and dry material are combined).

|  | Cytotype | N | Mean (pg) | Standard deviation | Min (pg) | Max (pg) |
| --- | --- | --- | --- | --- | --- | --- |
| Fresh material | a | 5 | 0.36 | <0.01 | 0.36 | 0.37 |
|  | b | 20 | 0.46 | 0.01 | 0.45 | 0.48 |
|  | c | 3 | 0.81 | 0.01 | 0.81 | 0.82 |
| Dry material | a* | 25 | 0.44 | 0.01 | 0.41 | 0.45 |
|  | b* | 21 | 0.54 | 0.01 | 0.52 | 0.57 |
|  | c* | 1 | 0.97 | -- | -- | -- |
| Fresh + dry material (*) | a+a* | 30 | 0.37 | 0.01 | 0.35 | 0.38 |
|  | b+b* | 41 | 0.46 | 0.01 | 0.44 | 0.48 |
|  | c+c* | 4 | 0.82 | 0.01 | 0.81 | 0.82 |
|  |  | 75 |  |  |  |  |

### Sex determination

All of the samples from SN group (29) and all the recombinant samples (15) were females, while the Ww group (38) consisted mainly of females and two males (one from Sierra Nevada Mountains and another one from the Alps), see Appendix 1. The high proportion of females in the Ww samples may be due to a collection bias, as we preferentially chose moss cushions with sporophytes, because sporophyte morphology is one of the few characters that enabled us to make a clear distinction in the field between *Ceratodon* and other morphologically similar genera (e.g., in the Bryales Limpr. and Pottiales M. Fleisch.). The presence of sporophytes, however, indicates that males must have been present. Male buds (perigonia) are deciduous, and may not be produced each season, meaning that sterile plants may have been males. In the Sierra Nevada Mountains we never observed sporophytes (the identity of all samples was verified in the laboratory by examining microscopic gametophytic characters). Similarly, Rams et al. (2014) reported finding no sporophytes in deep sampling carried out in Sierra Nevada Mountains from early spring to autumn from 2002 – 2004, and García-Zamora et al. (1998) found only one fructified specimen identified as *C. purpureus* in a survey of a zone close to Sierra Nevada in 1990-1991. Moreover, none of the *Ceratodon* samples from south eastern Spain in the MUB and GDA/GDAC (University of Granada, Spain) herbaria showed sporophytes. If we exclude a possible bias in the case of the Sierra Nevada Mountains samples, we can conclude based on the binominal distribution that the 95% CI for the probability of encountering males in the SN-type populations lies in the range of p = 0.00 for the lower limit and p = 0.12 for the upper limit for both tested methods (“Wilson” and “Clopper-Pearson”), which means that males might even be completely absent.

## DISCUSSION

In most major models of speciation, a period of allopatry is essential to evolve reproductive isolation (Coyne and Orr, 2004). However, in many cosmopolitan species, including many mosses and ferns, the entire habitable range of species is within the range of the dispersal distance of spores (Muñoz et al., 2004; Frahm, 2007; Pisa et al., 2013) making strict allopatry unlikely. Therefore, it is reasonable to propose that speciation mechanisms that either occur in sympatry or accommodate some gene flow contribute to generating the extant diversity in such groups. The two best-studied sympatric speciation mechanisms in plants are polyploidy and the evolution of self-fertilization (Barringer, 2007). Here we show that the evolution of a new species, closely related to the cosmopolitan *Ceratodon purpureus*, was associated with a 25% increase in genome size and a significant decrease in frequency of males (Nieto-Lugilde et al., under review). Surprisingly, although we have found neither males nor evidence of recent sexual reproduction (i.e., sporophytes) in the new species, the genetic diversity among members of this species is relatively high. Despite the long period of isolation suggested by the sequence divergence between *C. purpureus* and the new species, we have found evidence of interspecific hybridization, suggesting that the new species apparently has retained the capacity for sexual reproduction. In a separate paper we discuss the taxonomic implications of this discovery (Nieto-Lugilde et al., under review). Here we use genealogical and genome size data to make inferences regarding the genetic architecture of speciation, and the demographic parameters that permit such divergence.

Taxonomists have struggled with species delimitation in the genus *Ceratodon* since the description of the genus. Burley and Pritchard (1990) found references for nearly 50 specific or subspecific taxa within *Ceratodon*, but based on an extensive survey of herbarium specimens recognized only four species, *C. antarcticus* Cardot., *C. conicus* (Hampe) Lindb., *C. heterophyllus* Kindb., and *C. purpureus*, including three infraspecific taxa (subsp. *convolutus* (Reichardt) Burley, subsp. *purpureus,* and subsp. *stenocarpus* (Bruch & Schimp.) Dixon). Previous molecular population genetic analyses indicated that disjunct populations of *C. purpureus* were sometimes very closely related, clearly showing that long distance dispersal, even among continents, was frequent enough to erase any signal of strong population structure (McDaniel and Shaw, 2005). However, these data did not provide strong genealogical support either for or against the existence of distinct species other than *C. purpureus*. Subsequent classical genetic analyses showed that geographically and ecologically distant populations were partially reproductively isolated from one another (McDaniel et al., 2007, 2008), but these appeared to be somewhat porous reproductive barriers, and it was unclear that the populations represented different species.

McDaniel and Shaw (2005) did find some isolates of *C. purpureus* that were genetically distant from the more common haplotypes found in northern temperate regions. Here we found strong evidence that haplotypes which are distantly related to the typical *C. purpureus* haplotypes are locally abundant in the Sierra Nevada Mountains of southern Spain. We also found populations containing SN haplotypes and recombinants, together with some rare samples with exclusively the typical *C. purpureus* haplotypes. To evaluate the possibility that the segregation of these divergent haplotypes in the SN populations represents the retention of ancestral variation in the species (i.e., coalescent stochasticity causing incomplete lineage sorting) we generated coalescent simulations using *BEAST and P2C2M. These analyses showed that the divergence between these two haplotypic classes was too great to be explained by coalescent stochasticity. The fact that this polymorphism is found in all of the nuclear loci that we sampled, and is geographically concentrated to the Sierra Nevada region, suggests that balancing selection is also an unlikely explanation. Collectively these data suggest that the SN haplotypes comprise a rare species sister to and partially reproductively isolated from the cosmopolitan *C. purpureus*.

The default mode for the evolution of reproductive isolation is allopatric speciation. The sympatric occurrence of typical *C. purpureus* haplotypes and SN haplotypes, even at modest frequencies, contradicts the suggestion by McDaniel and Shaw (2005) that the Mediterranean populations were isolated from the rest of the species as a result of decreased spore rain in peripheral populations separated by prevailing global wind patterns. If we assume that the current dispersal capabilities of *C. purpureus* represent the ancestral condition, this suggests that geography may not have been the primary isolating mechanism between the nascent species. Morphological analysis of plants of both species grown in a common garden (as well as putative recombinants between them; Nieto-Lugilde et al., under review) indicate that members of Ww group can be distinguished morphologically from the SN group based on multivariate biometrical evaluation of microscopic features of the caulidia and phyllidia (stem length, presence or absence of apical comal tuft, leaf size and shape, leaf costa width at base of lamina and leaf costa excurrence). Nevertherless, we were unable to distinguish between the SN group plants and recombinants in field samples, suggesting that the environment influences the variance in taxonomically important characters. It is certainly possible that an extrinsic factor, like a habitat preference, isolated the two species.

It is also possible that an intrinsic factor isolated the two species. Remarkably, however, we detected only females in the SN species, suggesting that male lethality could contribute to isolating the two species. Sex in dioecious bryophytes like *C. purpureus* is determined at meiosis, by the segregation of a UV chromosome pair, meaning that *∼*50% of the spores produced in a population should be males. Some meiotic sex ratio variation has been observed in this species in natural populations (overall mean of proportion of males was 0.41 (0.17–0.72), Norrell et al., 2014) and artificial crosses (male-biased sex ratio = 60%, McDaniel et al., 2008). Even given our sample size (n = 29, with no males), we can conclude that the percentage of males in the SN populations is much lower (95% CI included 0% - 12%; additional samples not included in this study lower the 95% CI to a range of 0% - 6.7%). We do not know whether the decrease of males coincided with the speciation event, or occurred subsequent to the evolution of reproductive isolation. The evolution of apomixis or obligate selfing from historically outcrossing lineages is a well-documented route to the evolution of new species in plants (Stebbins, 1974; Barrett, 2010; Wright et al., 2013), and parthenogenetic lineages associated with the loss of males are frequent in some animal lineages (insects: Hagimori et al., 2006 and Montelongo and Gómez-Zurita, 2015; vertebrates: Neaves and Baumann, 2011 and Gutekunst et al., 2018). However, we know of no other cases where the loss of males has been associated with speciation in bryophytes.

The presence of recombinants containing both typical *C. purpureus* alleles and alleles from the SN species indicated that rare interspecies hybridization has occurred between individuals of the two species. Most of the recombinants possessed the SN chloroplast type, based on the *trnL* sequence data, suggesting that this species was more often the maternal parent (consistent with the rarity of males). We found one instance of a recombinant plant that had a typical *C. purpureus trn*L sequence, but we cannot determine whether this was a rare case of a hybridization involving a SN male (i.e., a cross in the opposite direction) or whether this resulted from a backcross of a male recombinant to a typical *C. purpureus* female. Intrinsic genetic incompatibilities are often manifest as Dobzhansky-Muller interactions, which result in asymmetric introgression patterns at the causative loci (McDaniel et al., 2008) due to the death of incompatible multi-locus genotypes. Although we sampled only six loci across the genome, the recombinants did have a tendency to have the SN alleles at the *TBP* and *rpL23A* loci. We are currently examining the frequency of polymorphism across the genome of the SN and recombinant genotypes to distinguish among forms of extrinsic and intrinsic isolation between the SN and typical *C. purpureus* populations.

The flow cytometric data also showed that members of the SN species had a genome ∼25% larger than typical members of *C. purpureus*. It is possible that the speciation involved a whole genome duplication event followed by rapid genome reduction, the duplication of a large chromosomes (Inoue et al., 2015; Panchy et al., 2016), or the accumulation of transposable elements (TEs), which contribute to the extraordinary variation in genome size within even closely related species in angiosperms (Vitte and Bennetzen, 2006). Although the current data represent the most comprehensive sampling of variation in genome size in *Ceratodon*, we still lack cytological data to determinate if variation in nuclear DNA content is due to an increase in the size of chromosomes or by the increase of number of chromosomes. The variance in genome size is almost equal between the two groups, suggesting that the SN species is fixed for whatever loci underlie the genome size change. Additionally, recombinants between the two groups have the genome size of SN species, not an intermediate value, suggesting that the increase in genome size may come from a single genomic change, rather than many small changes across genome. One hypothesis is that these plants have gained DNA on the sex chromosome which comprises nearly one-third of the genome (Heitz, 1932; Jachimsky, 1935; McDaniel et al., 2007). Sex chromosomes in other organisms are known to accumulate genomic material rapidly, sometimes in large translocations, and potentially generating pronounced evolutionary and ecological consequences (Tennesse et al., 2017). We are now attempting to generate artificial crosses to evaluate the genetic basis of the genome size difference.

We also found a third rare cytotype with a genome size approximately twice that of either SN plants or typical *C. purpureus* plants. These isolates all had mixed haplotypes (i.e., gene sequences from both the SN and typical *C. purpureus* clades) and a genome size very close to the sum of the SN group and Ww group (∼1.2 % smaller than the sum of the group means), suggesting that they arose from an allopolyploid event. Without more sequence or cytological data we cannot formally eliminate the possibility that the larger cytotype arose from autopolyploidy followed by hybridization, although this would require the gain of ∼10 % or loss (∼12 %) of the genomic DNA. Additionally, allopolyploidy is a widely observed mechanism to restore the fertility of F1s hybrids between partially reproductively isolated species with karyotypic differences and exhibit meiotic abnormalities (De Storme and Mason, 2014).

Finally, the new SN species apparently maintains levels of genetic diversity nearly equivalent to typical populations of its sister species *C. purpureus* without obviously undergoing sexual reproduction. Sexual reproduction in mosses occurs when males and females grow in close proximity, and sperm cells disperse, typically in humid conditions, from male to female plants, producing sporophytes, a very common observation in most populations of *C. purpureus*. Given the complete absence of sporophytes in observed Sierra Nevada samples, the predominant reproduction way seems to be by fragmentation of the gametophores. Moss gametophores can persist for many years, even in relatively stressful conditions, and easily spread clonally by fragmentation. In some cases, such fragments may be dispersed a considerable distance (Frahm, 2007, Lewis et al., 2014b). It is clear that spatially heterogeneous selection (Vrijenhoek, 1978) or frequency-dependent selection (Weeks and Hoffmann, 2008) can maintain high genetic diversity in clonal organisms. Antarctic populations of *C. purpureus*, which similarly lack any sexual reproduction, were also quite variable, although less polymorphic than was observed in the closely related nearby populations from Australia (Clarke et al., 2009). Also similar to the Antarctic studies, we found polymorphic nuclear ITS sequences between samples collected a few meters apart (unpublished data), indicating that these localities were colonized several times independently. However, unlike the Antarctic case, the SN isolates are genetically distinct from any known spore source. It is possible that sexual reproduction in the SN species generated the current variation under a past climate regime, or in undetected localities, although it is clearly far rarer than in *C. purpureus*. Further analyses of the evolutionary history of the SN population are likely to produce a better understanding of the phenomena that generate new species in cosmopolitan taxa.

## ACKNOWLEDGEMENTS

The authors thank the curators of herbaria BOL, GDA/GDAC and S for providing plant material. To Laura Forrest from Royal Botanic Garden Edinburgh, United Kingdom, for donating samples. To the management group of the National Park of Sierra Nevada Natural Space for allowing us to collect samples in the National and Natural Parks. This study was supported financially by the Spanish Ministry of Science and innovation (Projects CGL2011-22936/BOS and CGL2014-52579-R) and FEDER funds of the E.U., and an NSF grant (DEB 1541005) to SFM. M N.-L. also thanks the Spanish Ministry of Science and innovation, for being received a “Formación de Personal Investigador” fellowship (FPI program 2012) (reference BES-2012-056799).

## Appendix 1.

Voucher information for the studied specimens. For each sequenced sample the next information is given: herbarium code; geographical origin, collection date (year-month-day), gender if known (F for female, M for male), presence of sporophyte if appropriate, indicated by an asterisk (*), GenBank accession numbers for the six loci studied, given in the next order: *hp23.9*, *PPR*, *rpL23A*, *TBP*, *TRc1b3.05* and *trn*L; sequences obtained by cloning are indicated by their GenBank accession number given in parentheses.

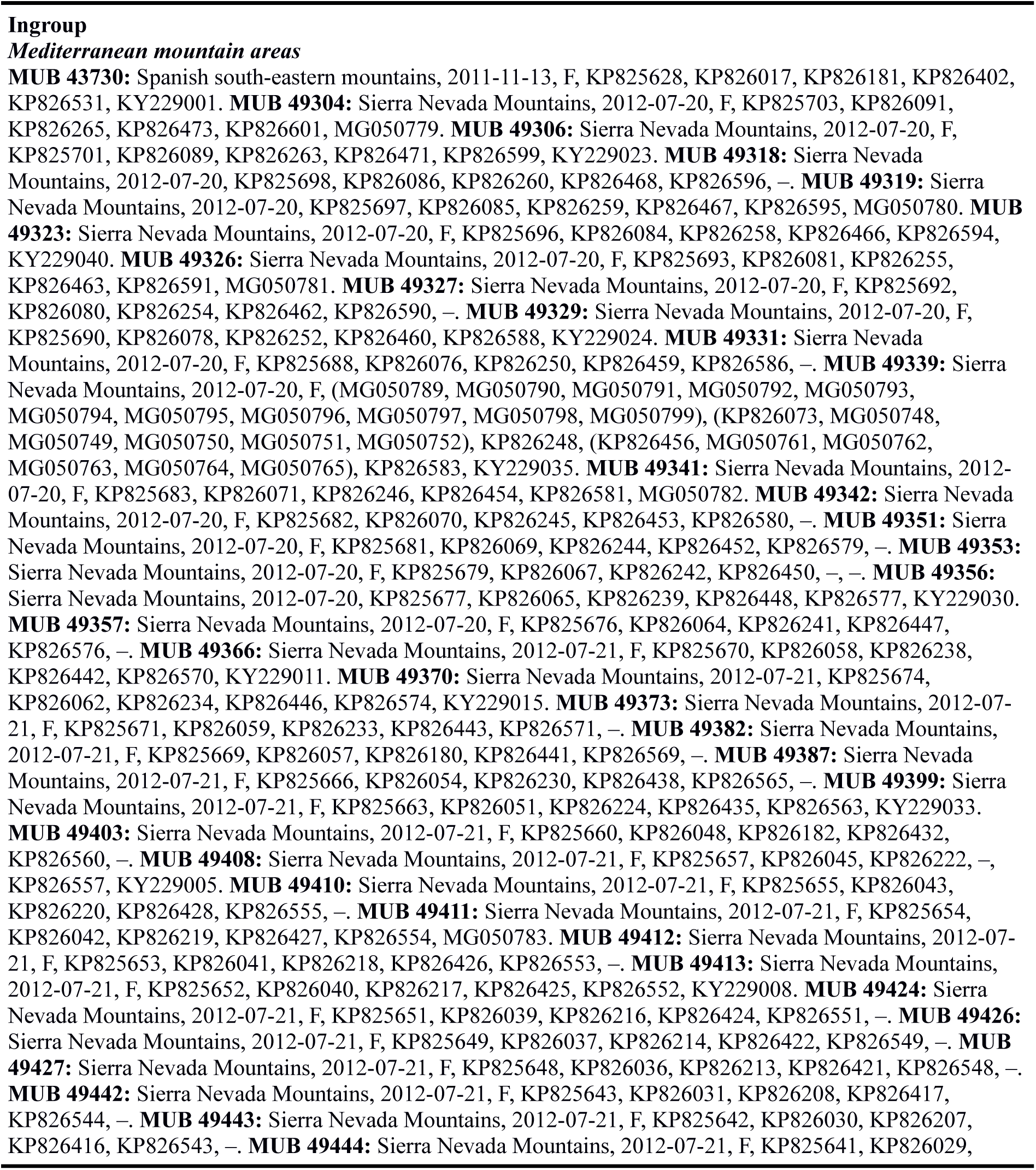
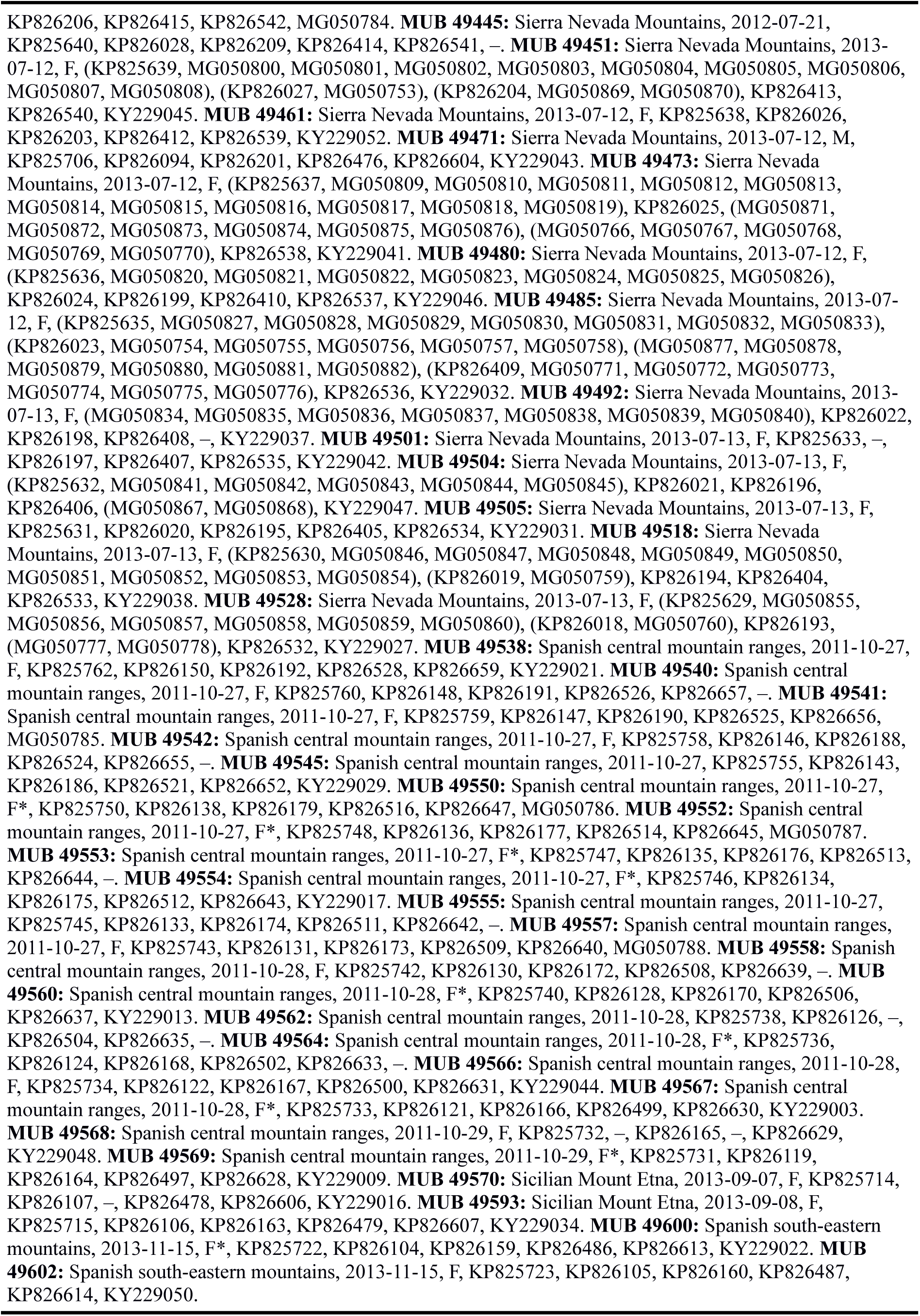
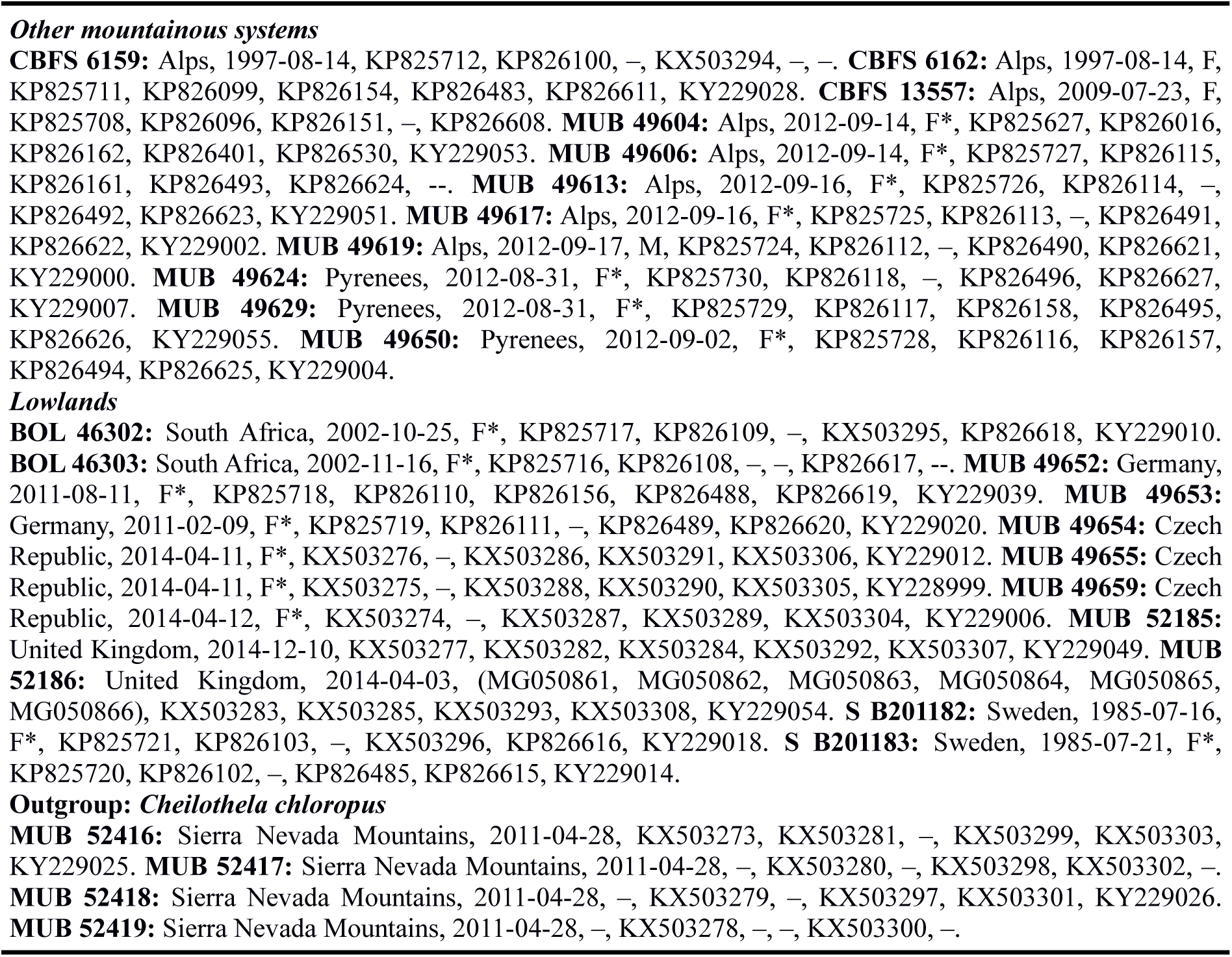

### Online Supplementary Materials

**Appendix S1.**
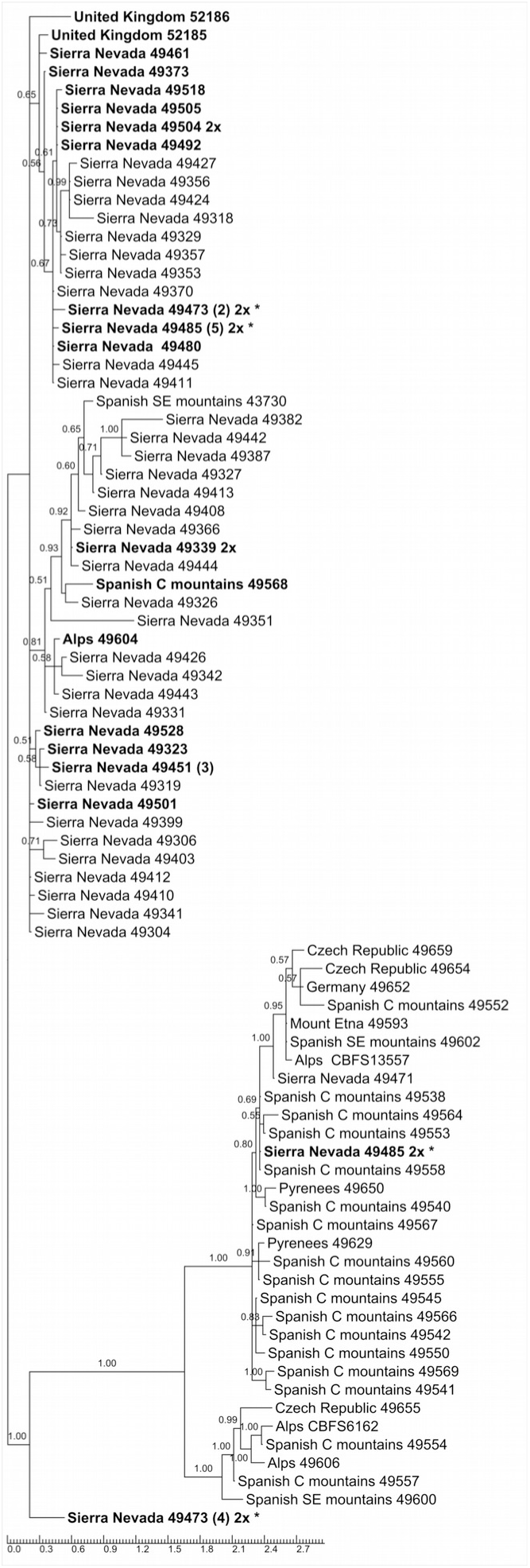
Phylogenetic tree inferred from the nuclear *rpL23A* locus. For each tip in the trees geographical origin and number of herbarium are given (numbers without letters are from MUB); 2x is used to highlight diploid samples; number of equal sequences obtained by cloning is indicated between parentheses if there was more than one; asterisk (*) is used for indicating samples with more than one copy for the locus; bold letters indicate recombinant samples.

**Appendix S2.**
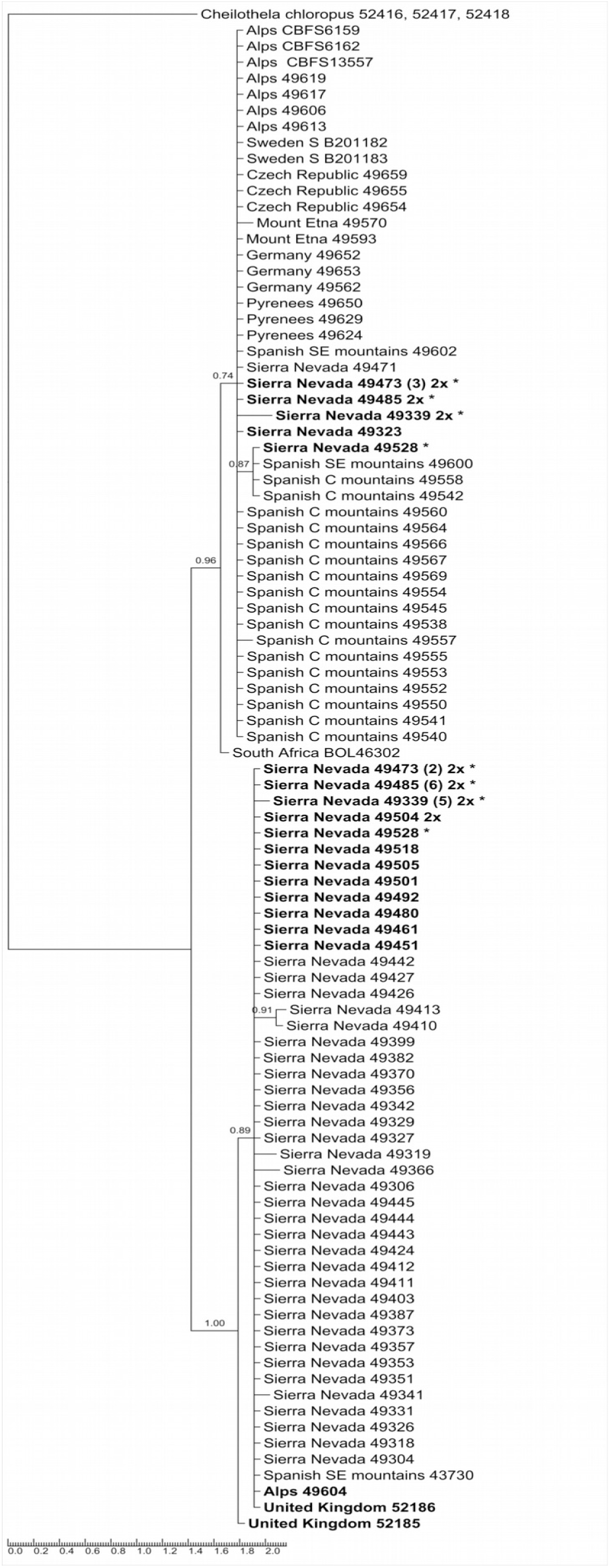
Phylogenetic tree inferred from the nuclear *TBP* locus. Information about the data given for each tip in the tree as in Appendix S1.

**Appendix S3.**
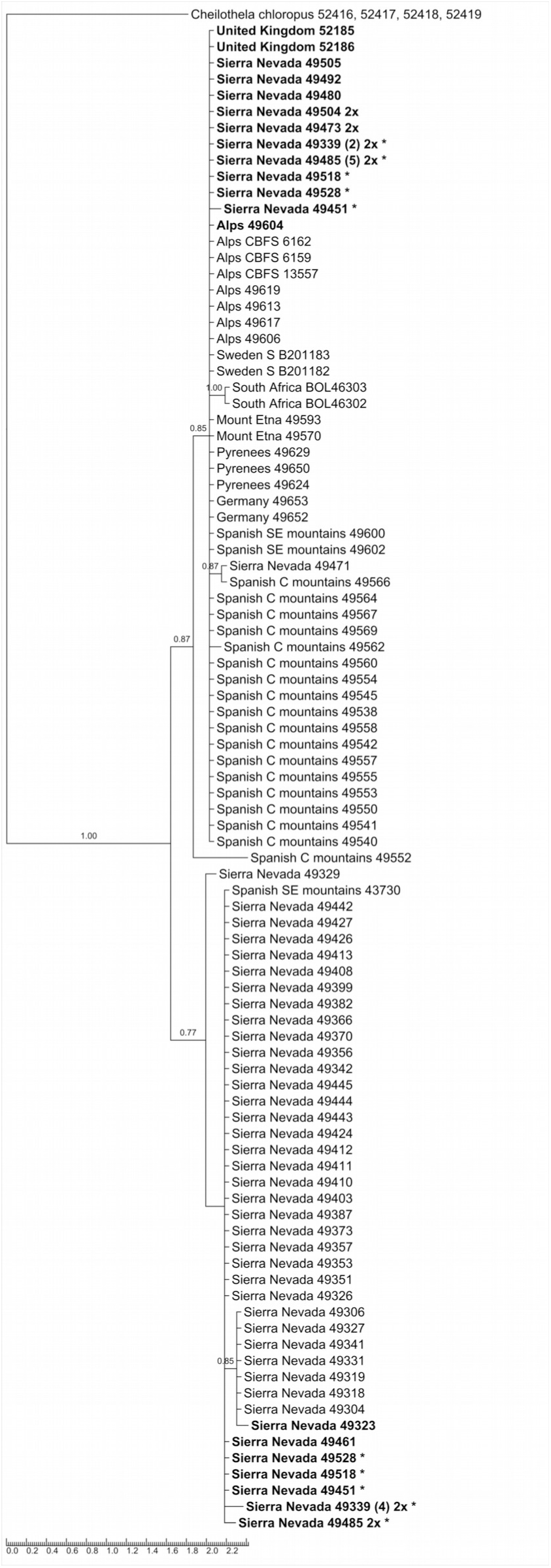
Phylogenetic tree inferred from the nuclear *PPR* locus. Information about the data given for each tip in the tree as in Appendix S1.

**Appendix S4.**
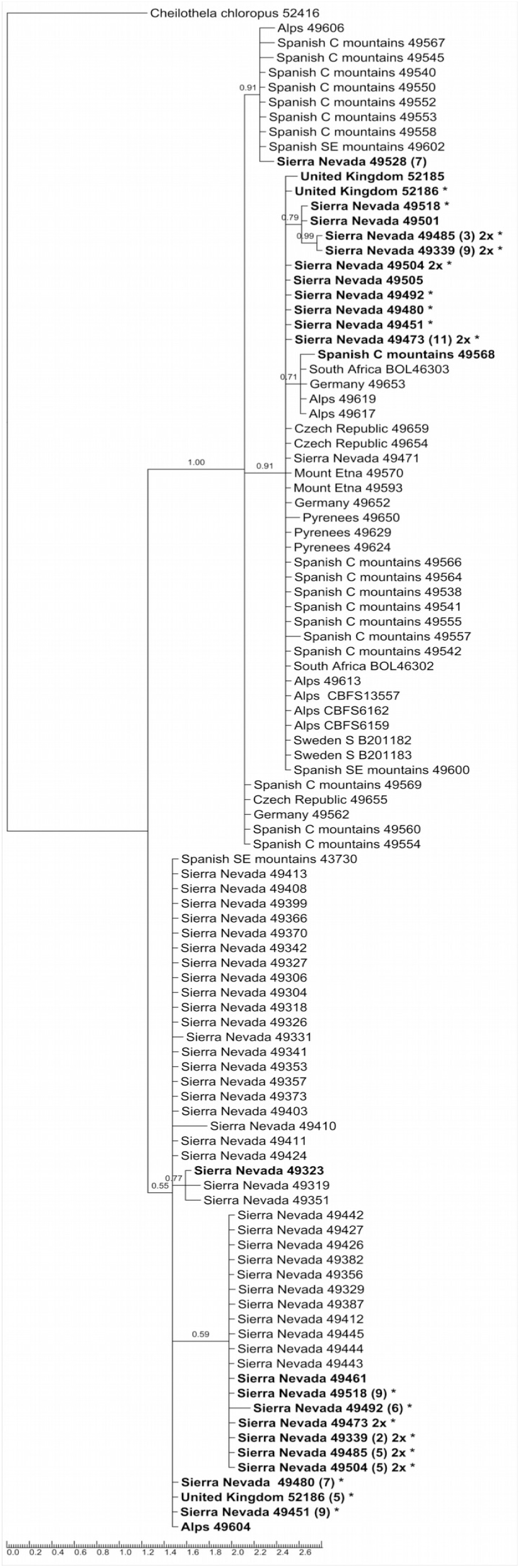
Phylogenetic tree inferred from the nuclear *hp23.9* locus. Information about the data given for each tip in the tree as in Appendix S1.

**Appendix S5.**
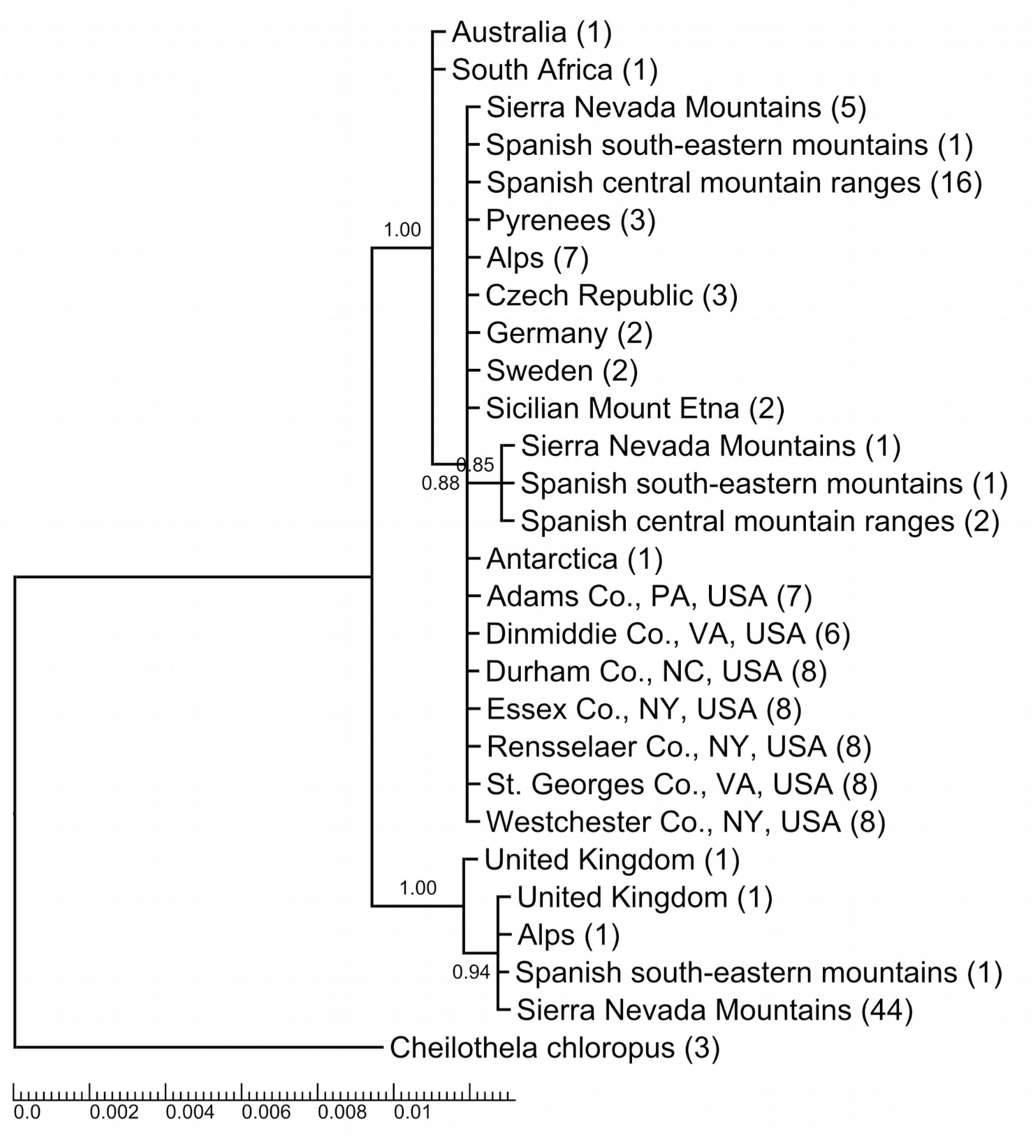
Phylogenetic tree inferred from the nuclear *TBP* locus adding to the samples used in this work other *Ceratodon* samples from Antarctica, Australia, and North America (GenBank accession numbers: KC436690 to KC436698, KC436701 to KC436706 and KC436710 to KC436750); To simplify the representation, we collapsed terminals of each group of individuals by area (total number of samples is given in parentheses).

**Appendix S6.**
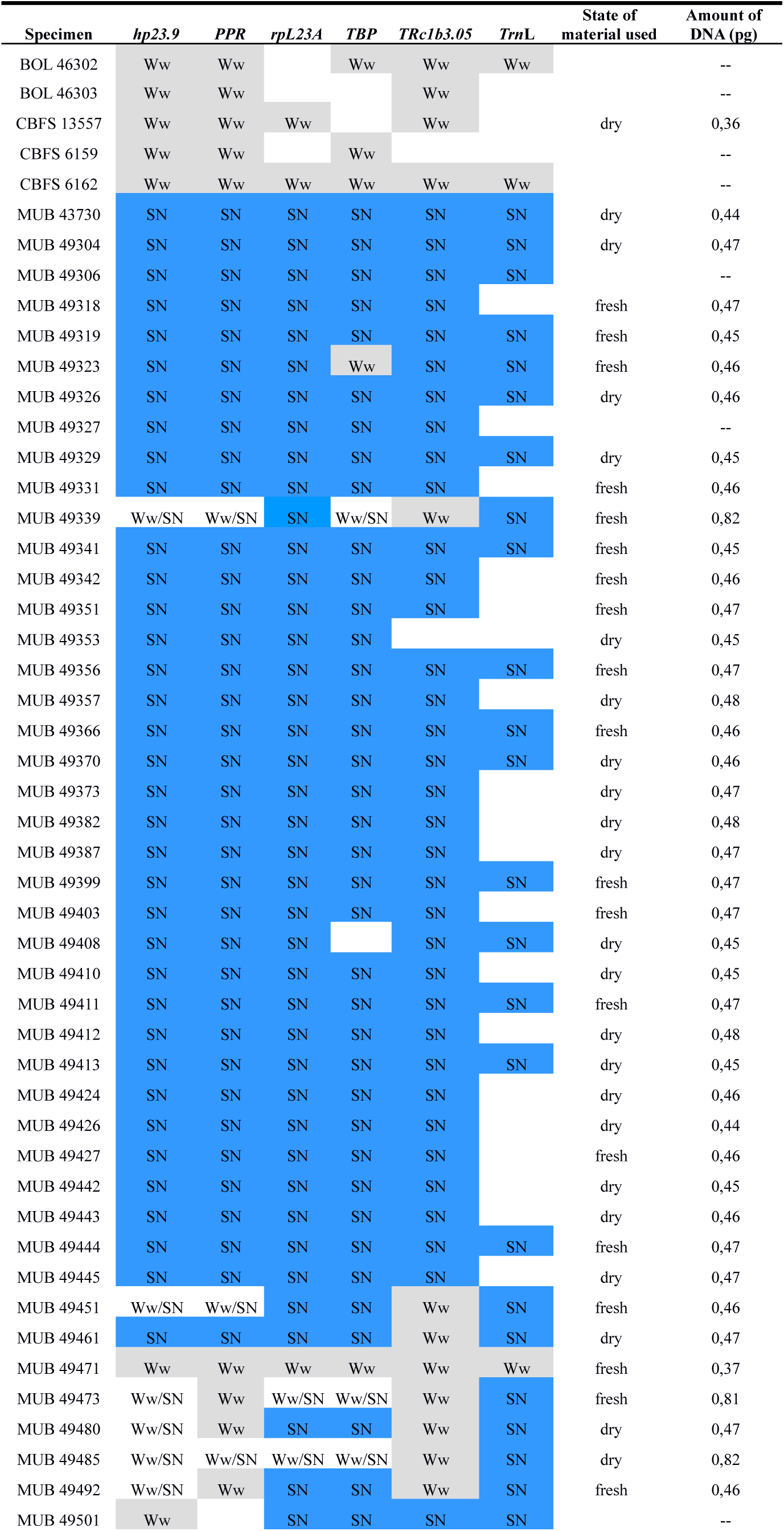
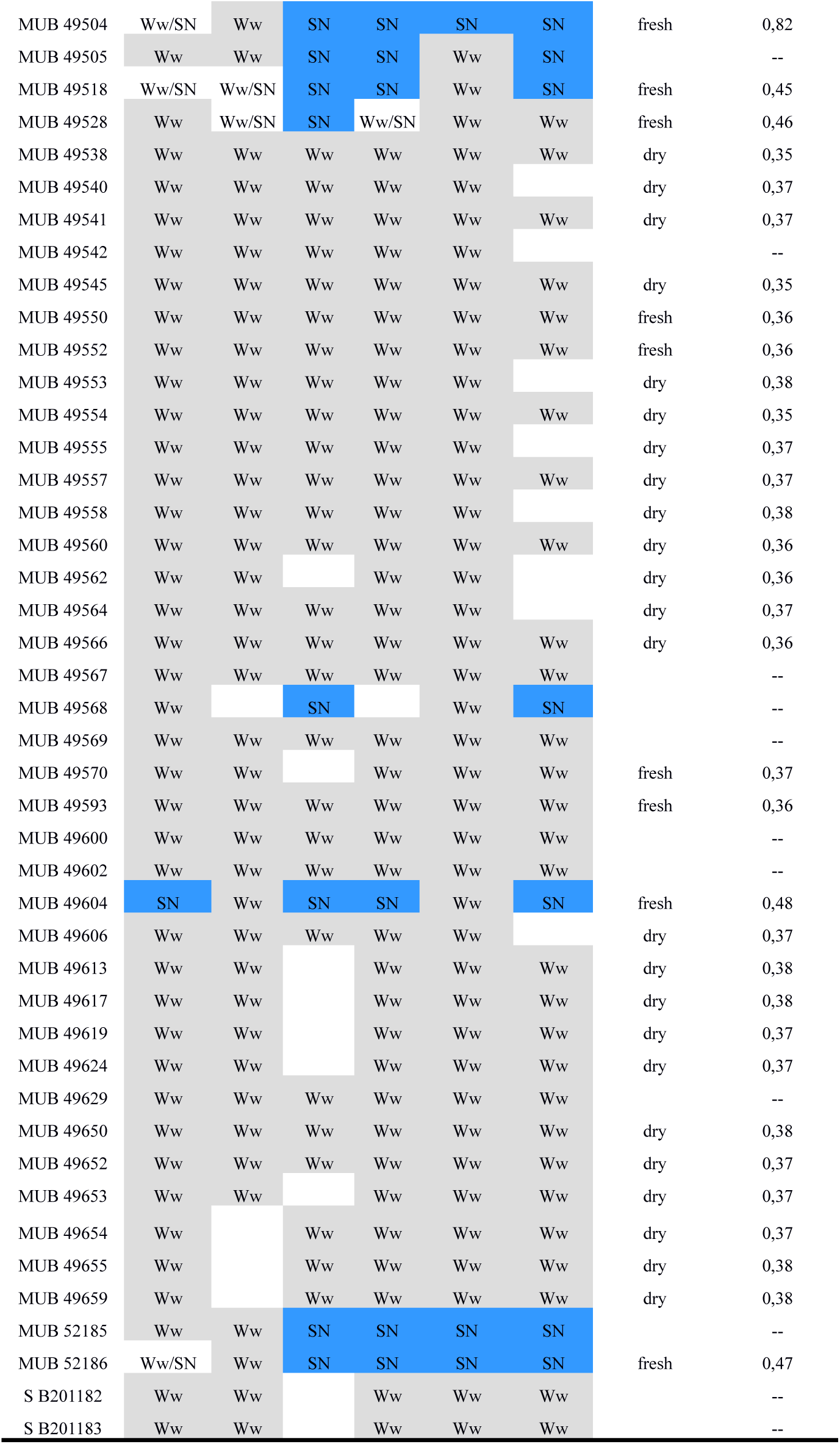
List of samples employed, indicating for each DNA locus analyzed, to which clade obtained in the phylogenetic analysis they belong (blue: SN clade, grey: Ww clade), the state of material used in cytometry analysis, and the amount of DNA (in case of dry material corrected by a factor of 0.85).

